# Heat tolerance and acclimation capacity in unrelated subterranean arthropods living under common and stable thermal conditions

**DOI:** 10.1101/598458

**Authors:** Susana Pallarés, Raquel Colado, Toni Pérez-Fernández, Thomas Wesener, Ignacio Ribera, David Sánchez-Fernández

## Abstract

Cave-dwelling ectotherms, which have evolved for millions of years under stable thermal conditions, could be expected to have adjusted their physiological limits to the narrow range of temperatures they experience and be highly vulnerable to global warming. However, the few existing studies on thermal physiology in subterranean invertebrates point that they have lost some of the thermoregulatory mechanisms common in surface species, but there is a lack of evolutionary adjustment to ambient temperature. The question remains whether this surprising homogeneous thermal niche is common for the whole subterranean biodiversity or just a peculiarity of the species tested. In order to test for the generality of such previous findings, we studied basal heat tolerance and thermal plasticity in four species of distant arthropod groups (Coleoptera, Diplopoda and Collembola) with different evolutionary histories but under similar selection pressures, as they have been exposed to the same constant environmental conditions for a long time (inhabiting the same cave). Adult specimens were exposed at different temperatures for one week to determine upper lethal temperatures. Then, surviving individuals from each treatment were exposed to a fixed stressful temperature to determine acclimation capacity. Upper lethal temperatures were similar to those reported for other subterranean species (between 20-25°C), except from that of the diplopod (28°C), widely exceeding the cave temperature (13-14°C). Acclimation responses differed from a positive acclimation response in two of the species to no acclimation capacity or detrimental acclimation effects in the others. Our results show that thermal niche features are not as homogeneous as expected across the subterranean fauna and unrelated to ambient conditions. We show that some species, likely those that colonized subterranean environments more recently, still retain thermoregulation capacity to face temperature changes. Thus, subterranean species, even those living under similar climatic conditions, might be very differently affected by global warming.

## Introduction

According with the climatic variability hypothesis [1], a positive relationship exists between the thermal tolerance and the level of climatic variability experienced by taxa, which has been demonstrated at different taxonomic resolutions (e.g. [2-5]). Similarly, thermal acclimation capacity is in general very limited in organisms from thermally stable environments compared with those living under more fluctuating conditions [6]. Thus, ectothermic animals which have evolved for millions of years under stable thermal conditions are generally highly stenothermal and lack physiological plasticity, which makes them especially vulnerable to global change [7-10]. For example, some Antarctic cold-stenothermal species (e.g. [7, 11-14]) as well as warm stenothermal coral reef fishes [15, 16] have lost the ability to activate a heat shock response via the expression of heat shock proteins.

The subterranean environment, characterized by extremely stable thermal conditions, provides another interesting system to test these general hypotheses of climatic variability [17, 18]. As a result of the strong selection pressures in these climatically buffered environments, cave-dwelling species might have adjusted physiological performance to the narrow range of temperatures they experience and therefore could be expected to be highly stenothermal compared with epigean taxa. Such thermal sensitivity, together with their generally poor dispersal capacity and limited possibilities for behavioural thermoregulation in a homogenous environment, would limit the potential of subterranean species to adapt to global warming [18, 19]. However, several decades ago, Vandel [20] questioned the stenothermal character of cave fauna, based on the little experimental data available at that time for some cave-adapted invertebrates, which showed that they generally survive long-term exposures at 20°C and can resist temperatures up to 25-29°C for short exposures (e.g. [21-24]). Mitchell [25] reported that the carabid *Rhadine subterranea* (Van Dyke, 1919) shows a seasonal shift in its temperature preference, showing some degree of thermal plasticity. Bull & Mitchell [26] demonstrated that the cave millipedes *Cambala speobia* (Chamberlin, 1952) and *Speodesmus bicornourus* Causey, 1959 could survive 30°C for short exposures (several hours) despite living at constant temperatures close to 20°C.

Experimental data on thermal physiology in subterranean species are still very scarce, which in part could be due to the difficulties in collecting large number of specimens required for experiments or rearing them in the laboratory. The few, more recent studies that have specifically measured thermal breadth and plasticity in subterranean species show different results depending on the trait measured. Some stygobitic (groundwater) microcrustacean species present the typical characteristics of stenothermal organisms, i.e., low physiological plasticity (e.g. [27]) and narrow thermal breadths of performance optimum (e.g. [28]), being even sensitive to changes in temperature 2°C below or above their habitat temperature [29]. In contrast, other closely related groundwater species have a much wider thermal ranges for performance and survival [28, 29]. Rizzo *et al*. [30] measured long-term (7 days) survival of several species of a clade of leiodid beetles at different temperatures. Tolerance ranges were much narrower than those of terrestrial insects living at the same latitude, but still wider than the temperatures experienced trough their evolutionary history and unrelated to their current habitat conditions. Furthermore, lethal thermal limits were very similar for all of the studied species irrespective of the climatic conditions of the areas where they live, and also similar to those observed in other leiodids [22, 31].

What seems to be common from all these studies is that most subterranean species, even those from relatively cold climates (e.g. [29, 32]) can survive temperatures around 20°C but not higher than 25-26°C in the long-term (> 7 days). Together, these results show that, in agreement with the climatic variability hypothesis, subterranean ectotherms have a lower heat tolerance than most epigean species, but also suggest a general lack of evolutionary adjustment to ambient temperatures (i.e., they are not strictly stenothermal). This would imply that species living in warmer areas are closer to their upper limits of thermal tolerance and therefore at higher extinction risk than those living in high altitudes (colder areas), contrary to the general predictions for surface species. Yet the question that remains is to what extent this apparently homogeneous thermal niche is common for the whole subterranean biodiversity. This question is difficult to be answered given the general lack of data on thermal physiology for the subterranean fauna. To fill pat of this gap, we studied heat tolerance, accounting for both basal (upper lethal temperature - ULT) and induced tolerance (plasticity of ULTs via acclimation), in four distantly related arthropod species representative of the invertebrate community of a single cave system (Cueva de la Murcielaguina de Hornos, SE Spain). They represent a set of species with different evolutionary origins but exposed to similar strong selection pressures (i.e., the same constant environmental conditions) for a long time.

## Material and methods

### Target species, sampling and holding conditions

The four studied species coexist in the same cave system, Murcielaguina de Hornos (Jaén, Spain), and belong to three different orders of arthropods. (1) *Glomeris* sp (Diplopoda: Glomerida: Glomeridae) is an undescribed species of a mega-diverse genus within the order Glomerida, and one of the few within the genus which is strictly subterranean [33]; it seems to be confined to some caves in south Spain (unpublished observations). (2) *Deuteraphorura silvaria* (Gisin, 1952) (Entognatha: Collembola: Onychiuridae) is a troglophile (i.e. not strictly subterranean) species distributed in Central and South Europe [34-35]. (3) *Speonemadus angusticollis* (Kraatz, 1870) (Coleoptera: Leiodidae) is an Iberian endemic species strictly hypogean in its southern distribution (Andalusia region) but endogean in the central part of the Iberian Peninsula [36], and (4) *Atheta subcavicola* (Coleoptera: Staphylinidae) is one of the few species of the genus *Atheta* associated with caves but also found in epigean environments (i.e., troglophile), distributed in the Iberian Peninsula and France [37-40].

None of the target species is included in national or international lists of protected or endangered species. The Murcielaguina de Hornos cave is located in the protected area “Parque Natural y Reserva de la Biosfera de las Sierras de Cazorla, Segura y Las Villas”. Field sampling was made with the corresponding permissions from the park authorities and the “Consejería de Medio Ambiente de la Junta de Andalucía”. Adult specimens of the four species were collected by hand, and found usually close to guano deposits. The temperature in the internal part of the cave (13-14°C) correlates with that of the mean annual temperature of the surface (13.9°C, Worldclim database, http://www.worldclim.org), as generally occurs in cave systems [18, 41, 42]. Specimens were transported to the laboratory under controlled conditions, with substrate from the cave in addition to moss to keep a high humidity. In the laboratory, the cultures were stored for two days before the experiments in closed plastic containers (10×15cm) with a white plaster layer (approx. 1cm) and the moss and cave substrate, at the approximate temperature of the cave (13°C) in an incubator (Radiber ERF-360). Specimens were fed *ad libitum* with freshly frozen *Drosophila melanogaster* throughout the entire duration of the experiments.

For the experiments, specimens of *Glomeris* sp., *A. subcavicola* and *S. angusticollis* were placed in the plastic boxes with the plaster substratum, 2 or 3 volcanic stones of a diameter of 1–3 cm and wet tissue paper. As *D. silvaria* specimens were not easily visible in such boxes given their small size and so were placed in Petri dishes also with a plaster substratum and a tissue paper saturated in water. The substratum, stones and papers were wetted daily, and trays with distilled water were placed in the incubator to keep a relative humidity >90%. Temperature and relative humidity inside the plastic boxes were recorded every 5 min with HOBO MX2301 dataloggers (Onset Computer Corporation, Bourne, MA, USA).

### Basal heat tolerance: upper lethal temperatures

To estimate species’ heat tolerance, we assessed survival at different temperatures during long-term exposure (7 days). The plastic boxes or Petri dishes with the specimens (N= 7–16 replicates per treatment) were placed in cabinets at 13 (control temperature), 20, 23, 25 or 30°C. Survival was checked every 24 h and specimens were recorded as alive if they were capable of some movement after a slight touch with a brush.

### Induced heat tolerance: acclimation capacity

Acclimation capacity of the four species was assessed using surviving specimens (N=5– 16 specimens per species and treatment) from the heat tolerance experiment (i.e., previously exposed at the different temperatures), which were transferred to a high fixed temperature. Such a fixed test temperature was set up considering the survival response at sublethal temperatures in the previous experiment: those species whose survival was higher than or close to 50% of the exposed specimens after 7 days at 25°C (*Glomeris* sp., *A. subcavicola* and *S. angusticollis*) were directly transferred to 30°C, while those with lower survival rates (*D. silvaria*) were transferred to 23°C. Survival was checked every 24 h for 7 days.

### Data analyses

All the analyses were performed in R v.3.3.3. To explore species’ basal heat tolerance, we used Kaplan–Meier survivorship curves [43] for comparison of survival at the different tested temperatures. Right censored data were specified for those individuals that were alive at the end of the experiment (see [44]). Survival data from each species at the end of the experiment (7 days) were fitted to a logistic regression model from which LT_50_ values were estimated using the *dose.p* function.

The effect of acclimation treatments on survival at a high fixed temperature was determined for each species by using GLMs assuming a Poisson distribution. Post-hoc tests with Bonferroni correction were used to compare pairs of treatments and specifically to test for positive acclimation responses (i.e. significantly higher survival in individuals from heat treatments than those of the control group) or detrimental acclimation effects (i.e. significantly lower survival in individuals from heat treatments than those of the control group). Additionally, we employed non-parametric tests (Kruskal-Wallis) and *kruskalmc* posthoc test [45, 46] for comparison with the previous, more conservative approach, and to test for the robustness of the results.

## Results

Raw data on survival time from both experiments are available in the Dataset S1.

### Absolute thermal tolerance

All the species showed ca. 100% survival at 13°C (control) and 20°C during the 7 exposure days, and a rapid mortality (between 24–48 hours) at 30°C. Therefore, survival at 23 and 25°C marked the differences in thermal tolerance among the species: *Glomeris* sp. showed the highest survival at these temperatures (100% survival for 7 days) and *D. silvaria* the lowest (i.e., rapid and high mortality in both treatments) (Fig 1). Mean LT_50_ values (7 days) ranged from 19.64±1.03 (*D. silvaria*), 24.98±0.52 (*A. subcavicola*), 25.18±0.75 (*S. angusticollis*) to 27.59±0.76°C (*Glomeris* sp.).

**Fig 1.**
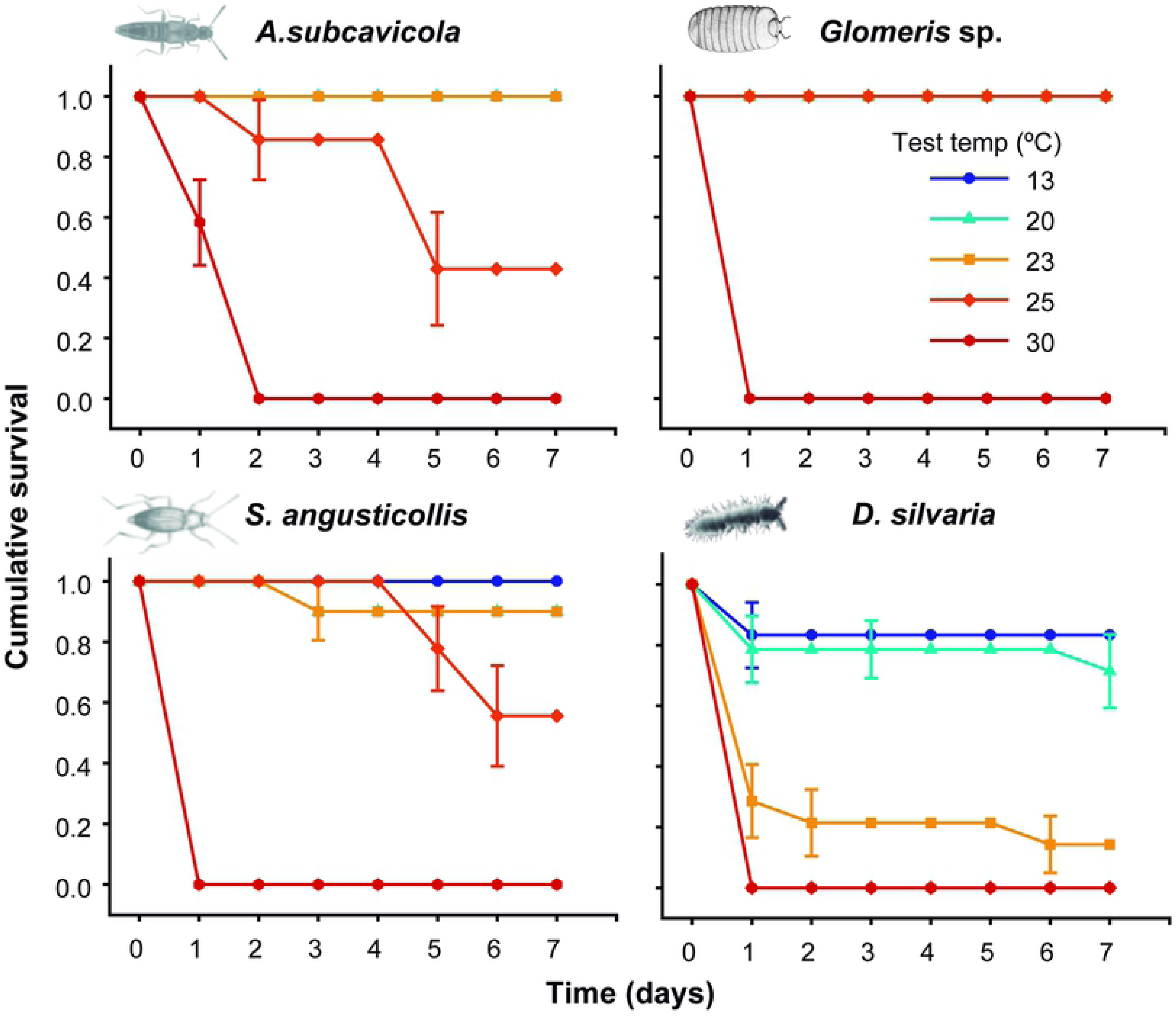
Kaplan–Meir survivorship curves for each temperature treatment. Each data point represents survival probability (mean±s.e.m.).

### Acclimation capacity

Each species showed a different response after acclimation at the different temperatures. *Glomeris* sp. and *A. subcavicola* showed a positive acclimation response, although in the latter only non-parametric tests detected significant differences among acclimation treatments (Table 1). In *Glomeris* sp., survival time at 30°C was significantly higher in individuals acclimated at 23 and 25°C than those from the control group, with a maximum difference of 1.8 days between 25°C and 13°C treatments. In *A. subcavicola,* only those individuals acclimated at 23°C showed a significantly higher survival time (1 day) than the control group (p<0.05 in *kruskalmc* post-hoc test), while no positive acclimation response was found at the other acclimation temperatures (Fig 2).

**Table 1.** Results of GLM and Kruskal-Wallis tests to determine the effect of acclimation temperature on the subsequent survival at a fixed temperature.

| Species | GLM (Poisson) |  |  | Kruskal-Wallis test |  |  |
| --- | --- | --- | --- | --- | --- | --- |
|  | X<sup>2</sup> value | df | p-value | X<sup>2</sup> value | df | p-value |
| *A. subcavicola* | 2.162 | 3 | 0.540 | 11.038 | 3 | 0.012 |
| *Glomeris* sp. | 14.496 | 3 | 0.002 | 40.137 | 3 | <0.001 |
| *S. angusticollis* | -- | -- | -- | -- | -- | -- |
| *D. silvaria* | 12.645 | 1 | <0.001 | 9.405 | 1 | 0.002 |

**Fig 2.**
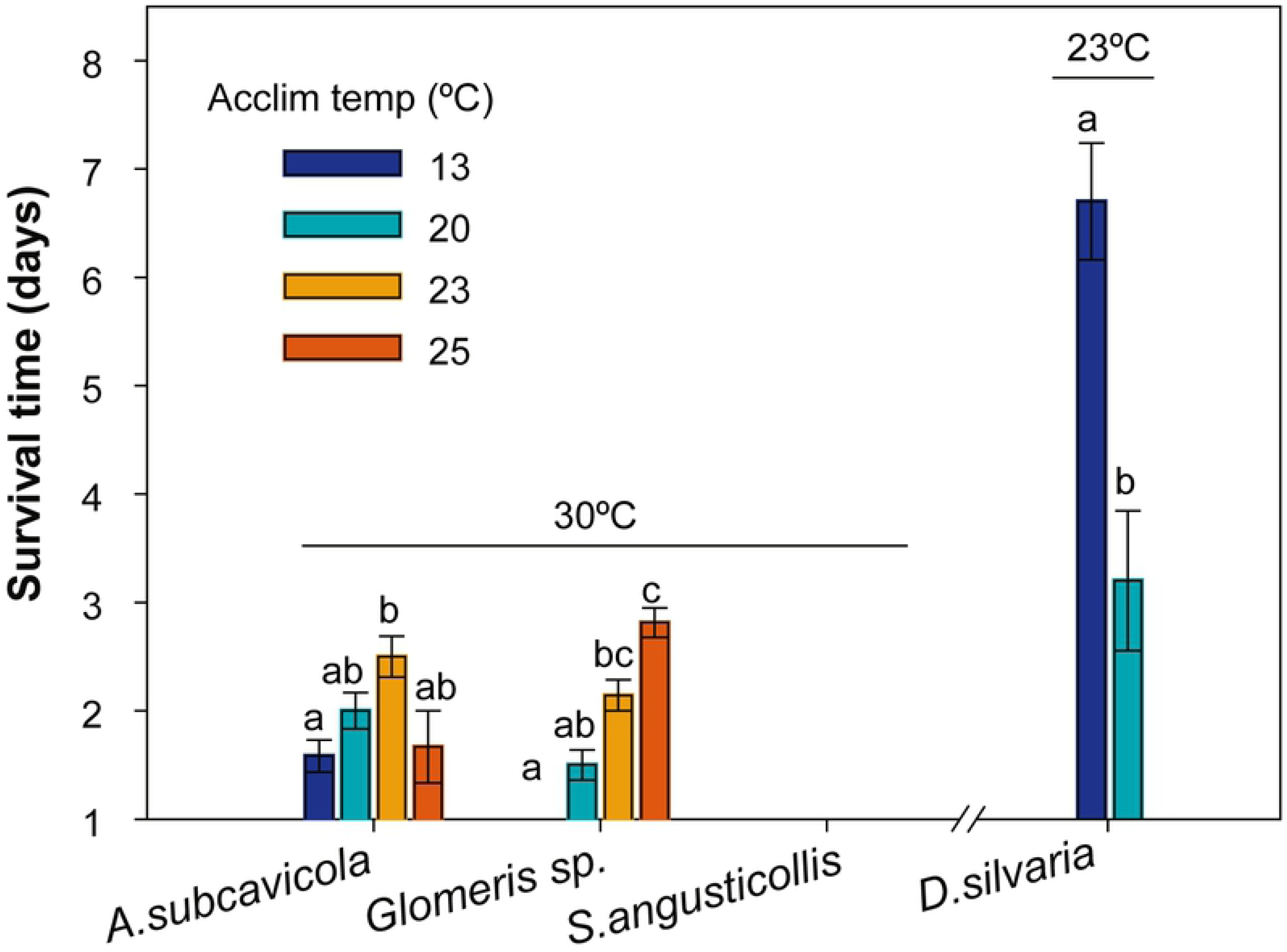
Survival time at a fixed temperature (indicated above each species) after acclimation at different temperatures. Letters above the bars indicate significant differences among acclimation treatments within species according to post-hoc tests (p<0.05).

*Speonemadus angusticollis* had no acclimation capacity: no specimens of any of the acclimation treatments survived more than 1 day of exposure at 30°C. In *D. silvaria,* acclimation at 20°C significantly reduced subsequent tolerance to 23°C (Fig 2).

## Discussion

As expected, and in agreement with the climatic variability hypothesis, upper thermal limits of the studied species were lower than those of most surface organisms [47]. However, the four species studied here can survive, at least for one week, at temperatures much higher than the narrow range currently experienced in their habitat (ca. 13-14°C). Our results are in agreement with previous work showing that subterranean species generally withstand temperatures up to 20°C with no apparent signs of stress (e.g. [28-30]. This contrasts with the narrow survival limits found in taxa from other thermally stable habitats, as for example stenothermal Antarctic fishes, which generally die at temperatures above 5°C [48], and most coldwater marine invertebrates, whose ULTs are generally not higher than 10°C [49]. Such higher thermal sensitivity of these marine species might be related with the role of oxygen in setting thermal tolerance, which is particularly relevant for aquatic organisms [50].

Nevertheless, thermal breadths for critical biological functions important to long-term survival or reproduction (e.g. feeding, locomotion, fecundity) may be more constrained than survival ranges in subterranean species (e.g. [29]). Similarly, exposure to non-lethal temperatures may have ‘hidden costs’ through sublethal effects (e.g. oxidative stress) which could affect fitness in the long-term [51]. It should be also noted that earlier stages could display different thermal sensitivity than adults. Indeed, data on the development of some strictly subterranean leiodid species showed high egg mortality between 15-21°C [31, 52].

We found some differences in survival at temperatures above 20°C with respect to other subterranean invertebrates. The two coleopteran species studied here showed LT_50_ values after 7 days exposure close to 25°C, a temperature which was lethal in brief exposures (<1 day) for several strictly subterranean species of leiodids [30]. But most notable was the high heat tolerance displayed by *Glomeris* sp., apparently the only troglobiont (i.e. obligate subterranean) species in our study, with a mean LT_50_ of 27.6°C. Although experimental data on thermal physiology are still scarce for subterranean ectotherms – and particularly for cave diplopods (but see [26]), to our knowledge, no known troglobiont species studied so far has shown such a high thermotolerance at long term exposures. Other than absolute lethal limits, their degree of plasticity also differed among the studied species. In comparative terms, thermal plasticity ranged from high in *Glomeris* sp. to moderate in *A. subcavicola* and absent in *S. angusticollis* and *D. silvaria*. In the light of these different acclimation responses, physiological plasticity deserves further exploration in subterranean species. Caves are environments with extreme selection pressures and adapted faunas that are similar in appearance, physiology, and behaviour all over the world, despite their different origins [53-55]. However, our results suggest that upper thermal limits and acclimation capacities, despite to be highly constrained, could be not as homogeneous as expected across the subterranean fauna. Instead they seem to be linked to particular aspects of the ecology, physiology or evolutionary history of the different lineages.

The degree of specialization to the subterranean environment, as well as habitat selection within the cave (both aspects related with the thermal variability to which organisms are exposed) have been associated with differences in the thermal tolerance among subterranean species. Some troglobionts have been shown to be less thermotolerant than facultative cave-inhabitants (trogloxenes and troglophiles) [56]. Such relationship was also consistent across a set of 37 phylogenetically distant invertebrate species [57]. Within troglobionts, species confined to the internal parts of the cave show also higher thermal sensitivity than closely related ones found close to cave entrances, where thermal conditions are more fluctuating [32]. Our small set of unrelated species prevents us to test this climatic variability hypothesis within an appropriate phylogenetic context. However, it is remarkable that, in contrast with these previous studies, the highest heat tolerance in our study was displayed by a troglobiont species which is found in the internal part of the cave Murcielaguina de Hornos, while the cohabitant troglophiles *A. subcavicola*, and especially, *D. silvaria,* were more heat-susceptible. Anyway, it would be interesting to explore intraspecific differences on thermal niche features related with the degree of specialization to the subterranean environment in these troglophile species, as their habitat preferences might change across their distribution range.

The differences in thermal physiological limits observed here between the studied species might thus reflect the particular evolutionary history of each lineage rather than distinct environmental preferences related to the occupied habitat. If thermal tolerance breadths have been progressively reduced in the process of specialization to underground environments, one might expect that lineages which have been isolated in caves for a longer time would show the most modified thermal tolerance ranges (i.e. most reduced with respect to that of the closest epigean ancestor), assuming a paradigm of evolution of thermal physiology traits similar to that of time-correlated troglomorphies [58, 59]. Currently, we still lack both molecular and physiological data needed to track the evolutionary reduction of thermal tolerance in the process of colonization of subterranean habitats. However, our results with *Glomeris* sp. suggest that this species may still retain some heat tolerance from a relatively recent epigean ancestor. Unpublished barcoding data shows a close relationship of the cave species (ca. 10% uncorrected p-distance) to *Glomeris maerens* Attems, 1927, an ecologically flexible species often found in semi-open habitat such as Mediterranean shrub and evergreen forests, at sea levels up to 1700 m elevation [60]. Despite showing other well-developed troglomorphic traits (depigmentation, reduced ocelli, elongation of body and appendages, see [54]), this species might belong to a lineage which colonized caves later than the other, less troglomorphic studied species. This also raises another interesting question for future research, as the asynchronous evolution of morphological and physiological traits in subterranean fauna.

Accurate predictions of species responses to climate change are mandatory if we aim to develop accurate management strategies to face this problem [9, 61]. In summer, maximum surface temperatures in the study site exceed or are very close to the ULTs of the species studied here. Although warming will be buffered and delayed in caves [62], these systems will not escape from the effects of climate change. We show here that subterranean species, even those living under the same climatic conditions, might be very differently affected by global warming. Among the studied species, the population of the collembolan *D. silvaria* in the Murcielaguina cave, in the margin and warmest extreme of its distribution range [34, 35], might be particularly threatened considering its low heat tolerance. Our results also stress the need of experimental approaches to assess the capability of species to cope with temperatures outside those they currently experience.

## Conclusions

According to the climatic variability hypothesis and similarly to other subterranean ectotherms, the species studied here showed narrow thermal tolerance ranges if compared with most terrestrial invertebrates. However, the estimated upper lethal limits largely exceed the current range of habitat temperature of these species. We also demonstrate that subterranean species with different evolutionary origins differ in their tolerance to heat and acclimation capacity, despite having been exposed to similar selection pressures (i.e., the same constant environmental conditions) for a long time. Therefore, thermal niche features of cave-dwelling species appear to be linked to particular aspects of the evolutionary history of the lineages (e.g. time of isolation in caves). Experimental data on thermal tolerance are essential for assessing the effects of global warming on subterranean fauna.

## Acknowledgements

We would like to thank the “Consejería de Medio Ambiente de la Junta de Andalucía” and “Parque Natural y Reserva de la Biosfera de las Sierras de Cazorla, Segura y Las Villas” for authorizing sampling at Cueva de la Murcielaguina de Hornos. We also thank members of the espeologist group of Villacarrillo (G.E.V) for conducting field work.

## Supporting information

**S1 Dataset. Raw data on survival time in basal heat tolerance and acclimation capacity experiments.**

